# N-VEGF, the autoregulatory arm of VEGF-A

**DOI:** 10.1101/2021.07.25.453721

**Authors:** Marina Katsman, Aviva Azriel, Guy Horev, Yitzhak Reizel, Ben-Zion Levi

## Abstract

Vascular endothelial growth factor A (VEGF-A) is a secreted protein that stimulates angiogenesis in response to hypoxia. Under hypoxic conditions, a non-canonical long isoform called L-VEGF is concomitantly expressed with VEGF-A. Once translated, L-VEGF it is proteolytically cleaved to generate N-VEGF and VEGF-A. Interestingly, while VEGF-A is secreted and affects the surrounding cells, N-VEGF is mobilized to the nucleus. This suggests that N-VEGF participates in transcriptional response to hypoxia. In this study, we performed a series of complementary experiments to examine the functional role of N-VEGF. Strikingly, we found that the mere expression of N-VEGF followed by its hypoxia-independent mobilization to the nucleus was sufficient to induce key genes associated with angiogenesis, such as Hif1α, VEGF-A isoforms, as well as genes associated with cell survival under hypoxia. Complementarily, when N-VEGF was genetically depleted, key hypoxia-induced genes were downregulated and cells were significantly susceptible to hypoxia-mediated apoptosis. This is the first reports of N-VEGF serving as an autoregulatory arm of VEGF-A. Further experiments will be needed to determine the role of N-VEGF in cancer and embryogenesis.

## Introduction

VEGF-A (or VEGF) is a key regulator of angiogenesis in normal physiology as well as in pathological events such as cancer (for review see [1, 2]). When translated under hypoxic conditions, a longer isoform of VEGF (L-VEGF) is generated from a non-canonical start site, CUG. L-VEGF adds 180 aa to the VEGF coding sequence and has been detected in VEGF-producing cells, including various cancer cells [3-5]. Once translated, L-VEGF is subjected to a proteolytic cleavage upstream to the classical translation start site of VEGF, yielding N-VEGF and VEGF (see schematic illustration in Fig. 1A). While VEGF-A is known to dramatically affect angiogenesis in surrounding cells the functional role of N-VEGF is not known. Interestingly, ectopically expressed N-VEGF shuttles to cell nuclei only under hypoxic conditions [6]. Furthermore, immunohistochemical staining of normal angiogenic tissue samples, such as kidney and breast, demonstrated sporadic staining of N-VEGF in the nuclei of renal tubular epithelium and of epithelium cells coating the mammary ducts. Finally, renal cell carcinoma and breast cancer cell nuclei exhibited strong immunostaining of N-VEGF in all cells [6]. These observations imply the involvement of N-VEGF in transcriptional regulation of VEGF-secreting cells. However, its actual impact on gene expression and on the response to hypoxia remains a mystery.

**Fig. 1.**
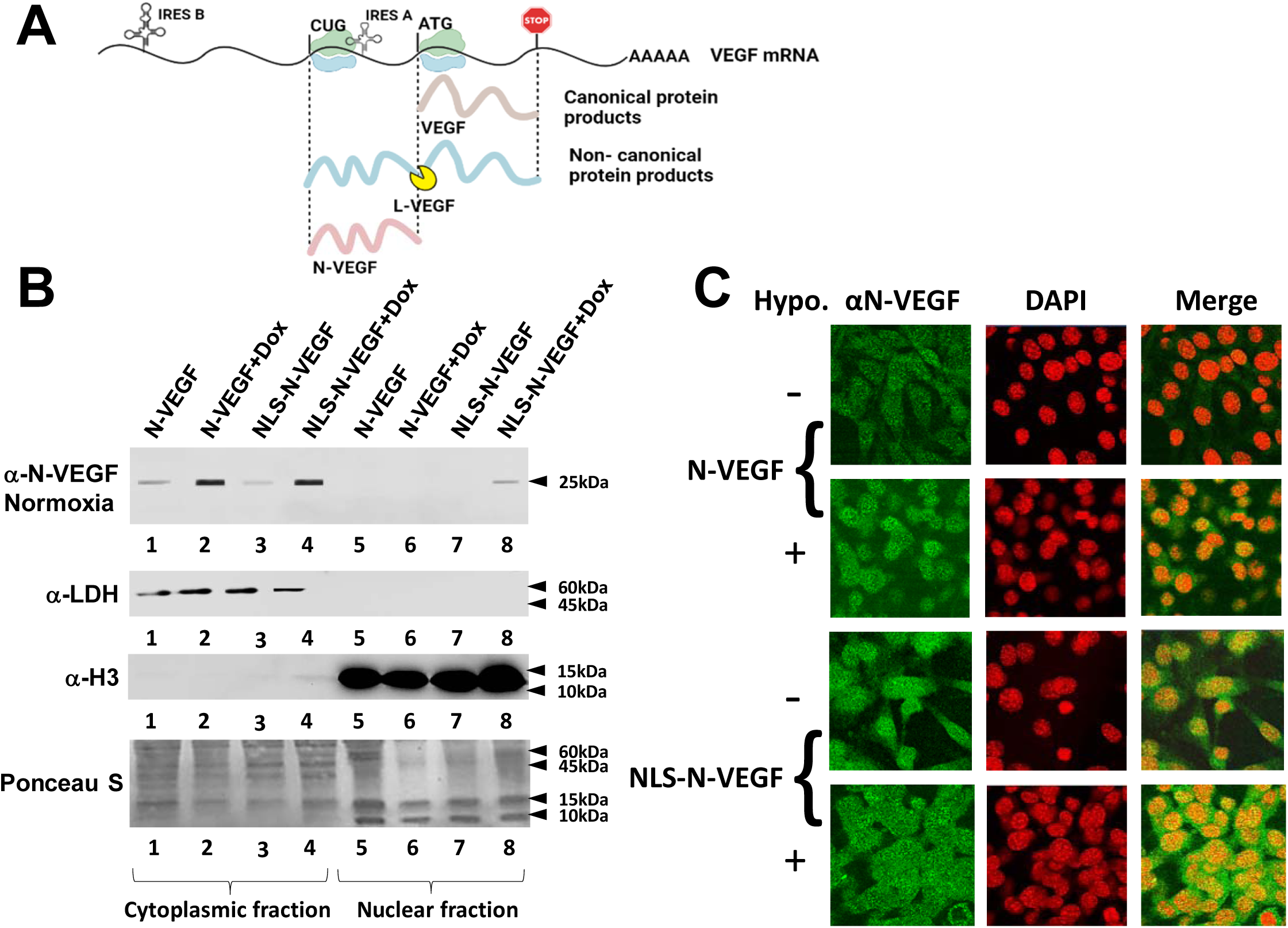
Hypoxia-independent nuclear mobilization of N-VEGF using the Cas9-NLS system. **(A)** Schematic illustration of the VEGF mRNA transcript, IRES domains and the translated products. Two translational products are generated. The non-canonical L-VEGF translation start site is CUG, and the canonical start site of VEGF is ATG. The locations of IRES A and B promoting translation under hypoxic stress are indicated. VEGF and L-VEGF proteins are illustrated as well as the proteolytic cleavage site of L-VEGF resulting in N-VEGF. **(B)** NIH3T3 cells were transduced with either N-VEGF or NLS-N-VEGF expression constructs without or with the inducer of the Tet-on promoter (Dox). Nuclear as well as cytoplasmic fractions were obtained from these cells. Western blot with anti N-VEGF antibodies was performed on these extracts. Subsequent Western blots on the same membrane were done using antibodies against LDH and histone H3 to show the purity of cytoplasmic and nuclei fractions, respectively. Ponceau S staining was carried out to demonstrate equal protein loading for all cytoplasmic and nuclear fractions. **(C)** Immunofluorescence of N-VEGF and NLS-N-VEGF cells before and following hypoxia (green) and nuclear staining with DAPI (red) is demonstrated.

We hypothesize that N-VEGF and VEGF play complementary regulatory roles. While VEGF is secreted and mainly affects neighboring cells, N-VEGF is retained in the cell and shuttles to the nucleus where it regulates the hypoxic transcriptional response. To characterize the role of N-VEGF in regulating transcription and in the adaptation to hypoxic stress, we performed a series of experiments. N-VEGF was mobilized to the nucleus, independent of hypoxia. Strikingly, its expression under normoxia was sufficient to upregulate key genes involved in the response to hypoxia and in the induction of angiogenesis. Complementarily, genetic depletion of N-VEGF inhibited the upregulation of hypoxia-induced genes and increased the apoptosis rate upon hypoxia. All these are sound proof for the key role of N-VEGF in the response to hypoxia.

## Results

### Expression and hypoxia-independent mobilization of N-VEGF

To examine if the mere presence N-VEGF in the nucleus is sufficient to induce a hypoxia-associated transcriptional response, we fused N-VEGF at its N-terminus to the nuclear localization sequence (NLS) of the Cas9 nuclease (NLS-N-VEGF). We chose to introduce N-VEGF to the nucleus under normoxic conditions to avoid the notable change that occurs in the transcriptome upon hypoxia [7]. For this purpose, N-VEGF and NLS-N-VEGF were cloned under the inducible Tet-on promoter in a lentiviral vector [8]. Subsequently, NIH3T3 cells expressing N-VEGF or NLS-N-VEGF, were grown in the presence or absence of the Tet-on promoter inducer, doxycycline (dox). NIH3T3 cells are mouse fibroblasts cells shown in many studies to activate hypoxia-associated genes, including VEGF-A and Hif1α, upon exposure to hypoxic stress [9, 10]. As seen in Fig. 1B, in the presence of dox, 25% of NLS-N-VEGF was identified in the nuclear fraction. The reason for the partial nuclear localization is probably because Cas9-NLS fusion to N-VEGF resulted in reduced translocation efficiency. It is important to note that under hypoxia, only 50% of the induced N-VEGF shuttled to the nucleus (Fig. 3A). Thus, only a 50% reduction in its nuclear mobilization was noted on addition of the NLS tag. Further, immunofluorescence analysis confirmed that N-VEGF exhibited nuclear localization only following hypoxia (Fig. 1C, second row), while NLS-N-VEGF showed nuclear expression even without hypoxia (Fig. 1C, third and fourth rows).

### Hypoxia-independent nuclear mobilization of N-VEGF induces part of the hypoxic transcriptional program

To study the effect of nuclear NLS-N-VEGF on cellular transcription, RNA was extracted from cells expressing either N-VEGF or NLS-N-VEGF and subjected to RNA-Seq. In efforts to identify altered gene expression resulting from shuttling of N-VEGF to the nucleus, while avoiding the transcriptional dynamics affected by the mere expression of N-VEGF in the cytoplasm, we only monitored genes exhibiting differential expression in dox-induced NLS-N-VEGF as compared to N-VEGF expressing cells and their dox untreated counterparts. Cells were collected after a 16-h treatment with dox to allow the translation and mobilization of N-VEGF to the nucleus. The differentially expressed genes were divided to up- and downregulated groups (fold change ≥ 2, adjusted p-value<0.05) and each group was functionally annotated using Metascape [11]. 123 upregulated and 32 downregulated differentially expressed genes resulting from the shuttling of N-VEGF to the nuclei were identified (see genes list and annotations in Table S1).

The annotated groups of the upregulated genes (Fig. 2A) clustered into three main categories. The first and most relevant category (Fig. 2A, purple bars), represent genes promoting angiogenesis, including Hif1α, a major hypoxia-induced transcriptional regulator [12], and VEGF isoforms 120 and 164 (Fig. 2B). Additional signature genes mediating angiogenesis, such as Adgrg1, Itgav, Nrp, Flna and Pik3c2a, were upregulated by mobilization of NLS-N-VEGF to the nucleus (Fig. 2B, and Table S1) [13]. Another dominant annotation involved genes associated with anti-apoptotic pathways rescuing cells from hypoxic stress (Fig. 2A, brown bars). Indeed, signature genes of anti-apoptotic pathways such as Sh3rf1 [14], Lrrk2 [15] and Src [16], were induced by N-VEGF shuttling (Fig. 2B, and Table S1). The last and major category of annotated gene groups was associated with integrity, structure, and maintenance of cellular cytoskeleton, which is highly relevant during hypoxic stress (Fig. 2A, red bars), including Cav2 [17], Lamc2 [18], Cfl2 [17], Rock2 [19], and more (Fig. 2B). Strikingly, many of the 123 upregulated genes are known targets of Hif1α, pointing to a key pathway activated by N-VEGF. Genes downregulated following NLS-N-VEGF mobilization to the nucleus included genes associated with regulation of translation, (Fig. S1). This aligns with the role of N-VEGF as an inducer of adaptive mechanism to hypoxic crisis by inhibiting the translational machinery [20].

**Fig. 2.**
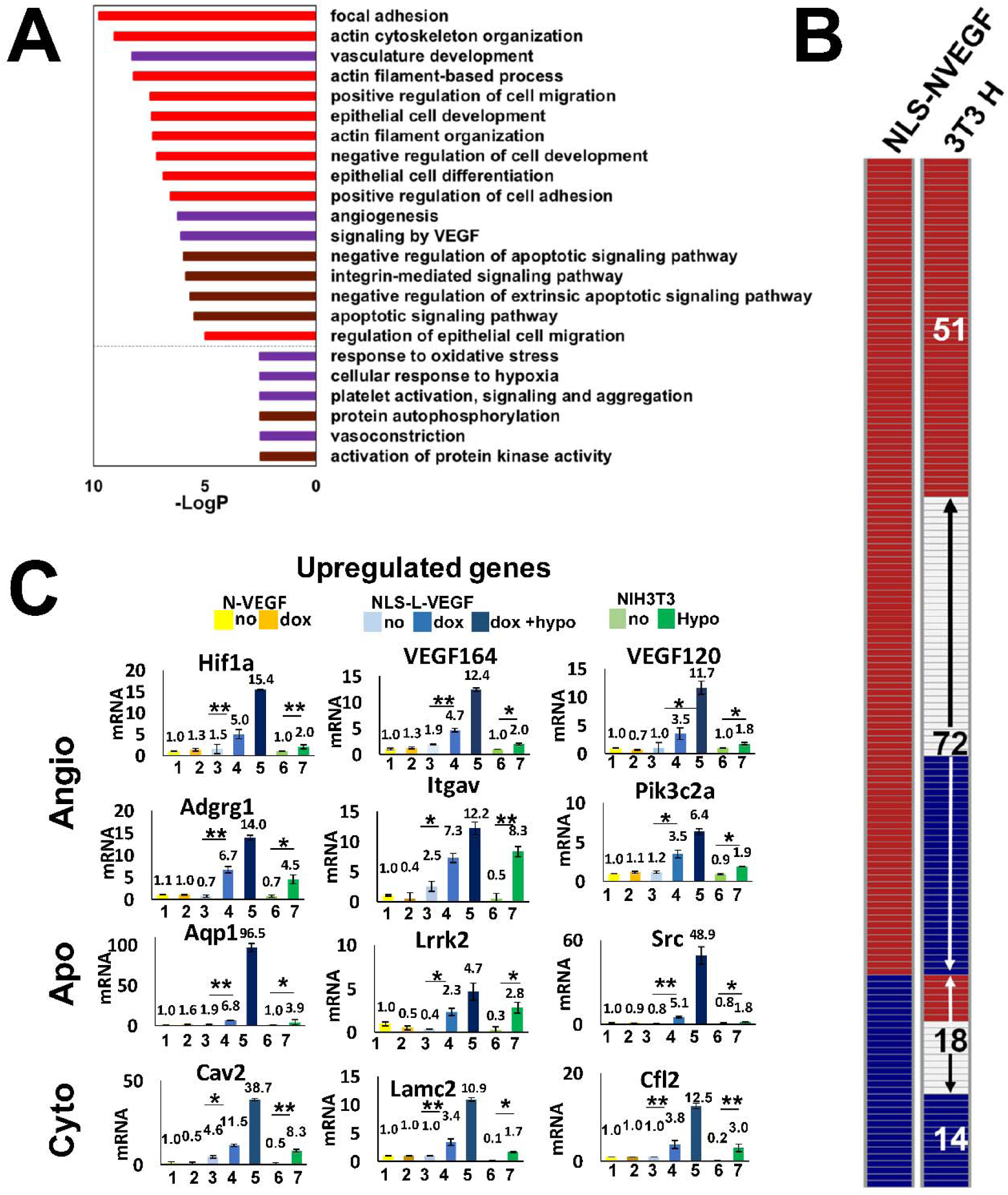
Transcriptional impact of hypoxia-independent mobilization of N-VEGF to the nucleus. **(A)** The top gene ontology terms of the 123 genes upregulated when N-VEGF is shuttled to the nucleus. **(B)** Qualitative comparison between 155 differentially expressed genes following mobilization of N-VEGF to the nucleus and their expression trend in wild type cells exposed to hypoxia. The columns represent a qualitative heatmap showing the direction of change in gene expression level (up-red, down-blue, no change-white) in NLS-N-VEGF cells following dox treatment (NLS-NVEGF, left column) in comparison to NIH3T3 cell under hypoxia (3T3 H, right column). The numbers within the right column represent the number of genes in each group that were annotated (detailed annotations in Fig. S2 A-D). **(C)** Validation of RNA seq results using qPCR. Real-time RT-qPCR analyses were performed on select upregulated genes from each functional annotation group. The data is presented for L-VEGF-expressing cells before (yellow) and following treatments with dox (dark yellow), NLS-L-VEGF cells before (light blue) and following treatments with dox (blue) and NLS-L-VEGF cells treated with dox plus hypoxia (dark blue). The data were compared to control NIH3T3 cells before (light green) and following hypoxia (green). Genes associated with angiogenesis (Angio), Apoptosis (Apo) and cell structure (Cyto) are indicated. Fold changes in mRNA levels in triplicates from three independent experiments were determined. Statistical significance was determined by two-way ANOVA, * p≤0.05, ** p≤0.01). Values are mean± AVEDEV.

**Fig. 3.**
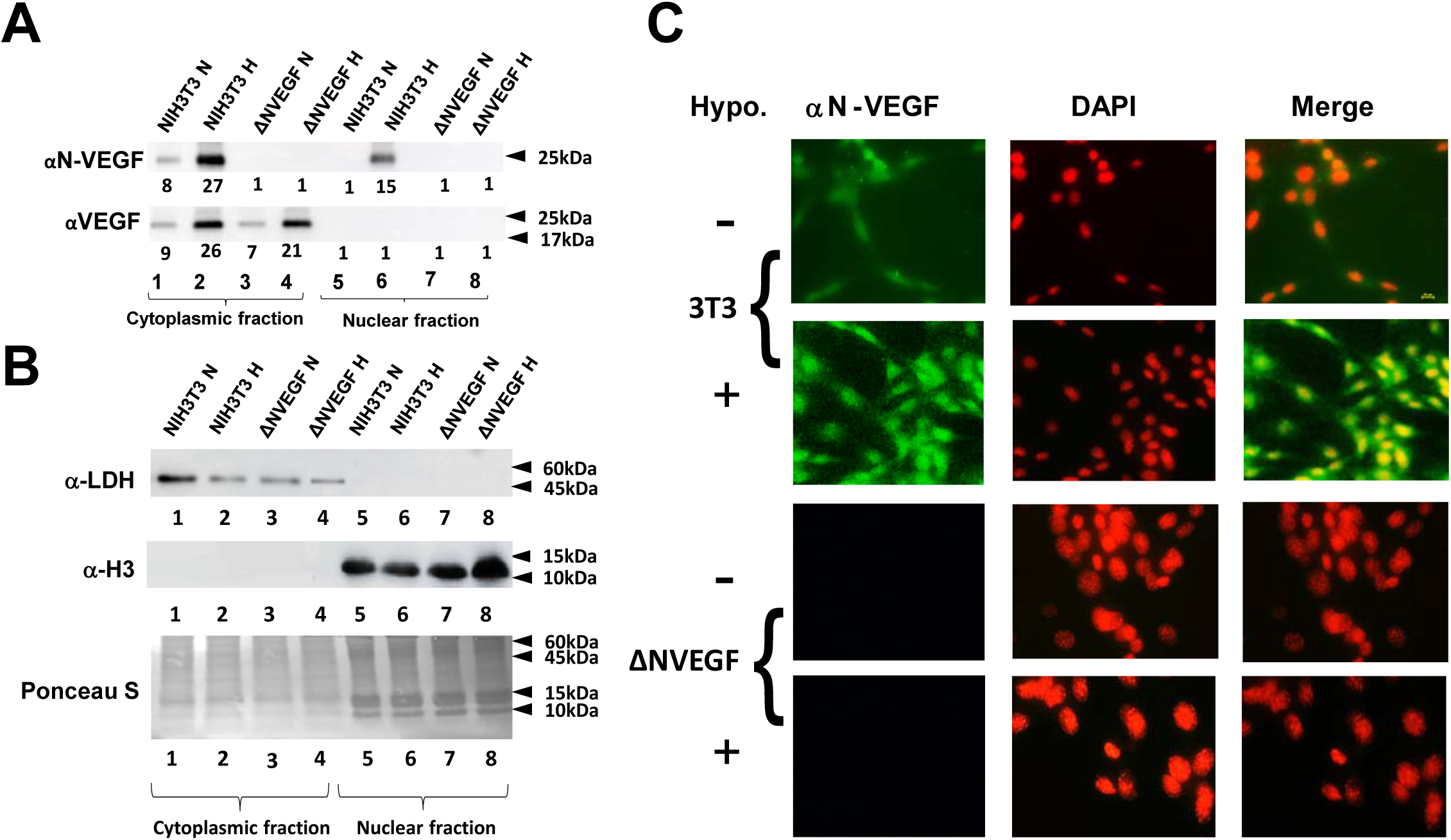
Genomic deletion of N-VEGF using CRISPR-Cas9. **(A)** Cytoplasmic and nuclear fractions were prepared from NIH3T3 and N-VEGF deleted cells (ΔNVEGF). Western blots were performed with antibodies directed against N-VEGF and VEGF showing complete deletion of endogenous N-VEGF using CRISPR-Cas9. Numbers below lanes indicate the average relative density of the band from three different gels as determined using ImageJ software [38]. Statistical significance was determined by Student’s t-test, p≤0.05. **(B)** To demonstrate the purity of soluble cytoplasmic and nuclear protein extracts, Western blots and Ponceau S staining were performed as elaborated in Fig.1 **(C)** Immunostaining showing complete absence of N-VEGF in deleted cells was performed as elaborated in Fig 1.

In order to examine whether the expression pattern of the 155 differentially expressed genes overlapped with genes induced by hypoxia, we compared N-VEGF-induced genes to hypoxia-mediated gene expression (elaborated hereafter, Fig. 4A). Overall, 42% of the genes exhibited similar trends, with 51 genes induced and 14 genes downregulated in both hypoxia-induced cells and in cells showing hypoxia-independent mobilization of N-VEGF (Fig. 2B). The annotated groups of upregulated genes that exhibited similar expression, grouped into the three main categories described above (Fig. S2A). These observations were verified by qPCR on a set of key genes in the three groups (Fig 2C). Exposure of NLS-N-VEGF-expressing cells to both dox and hypoxia resulted in a synergistic effect, with increased expression levels of these key genes (Fig. 2C, dark blue columns). Last, the critical role of nuclear mobilization of N-VEGF in regulating the expression of these genes was demonstrated by the lack of an effect of dox on N-VEGF-infected cells. Taken together, this focused gene profile expression analysis highlighted the role of nuclear N-VEGF in initiating part of the main hypoxic signaling cascade.

**Fig. 4.**
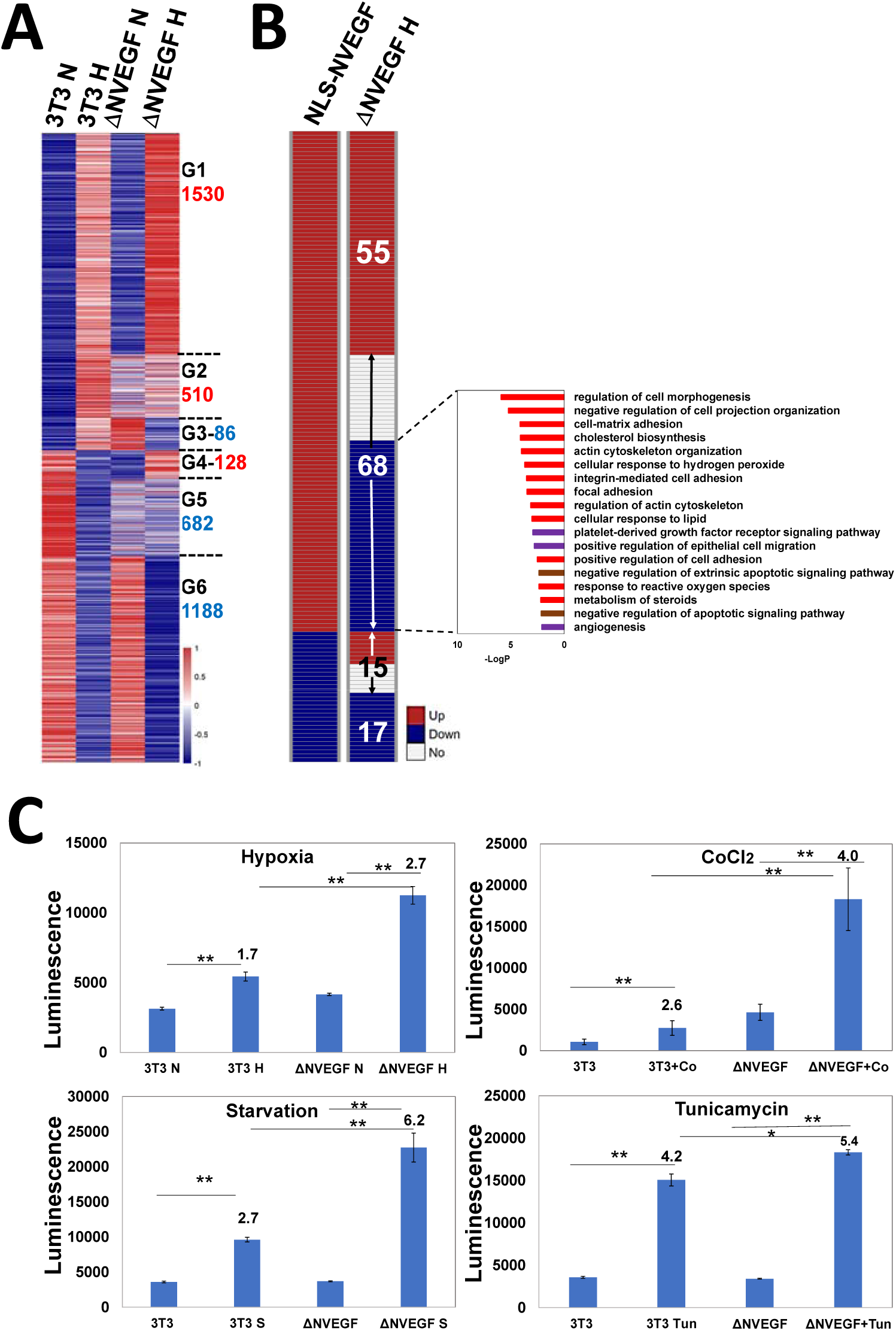
Impact of N-VEGF deletion on hypoxia-induced genes and on cellular physiology. **(A)** Shown is a heatmap demonstrating gene expression levels in NIH3T3 cells (3T3) and ΔN-VEGF counterpart cells (ΔNVEGF) under normoxia (N) and following hypoxia (H). The heatmap presents only the 3,000 genes modified following hypoxia in the wild type. The genes were divided into six groups (G1-G6) of upregulated (red), downregulated (blue) expression in hypoxia. The numbers of differential genes in each group is indicated next to the name of each group (red-up and blue-down). **(B)** A qualitative comparison between the 155 genes that exhibited differential expression when N-VEGF was shuttled to the nucleus and the expression trend of these genes in ΔN-VEGF cells following hypoxia. The numbers of genes in each group is indicated in the right column. The annotations of genes (68) that were upregulated in NLS-N-VEGF cells following dox and downregulated in ΔN-VEGF cells are presented. **(C)** To induce stress signals, NIH3T3 cells and their N-VEGF-deleted counterparts were exposed to hypoxia, CoCl2, starvation, and tunicamycin for 18 h. Apoptosis was determined using the ApoTox-Glo™ Triplex Assay. The results represent three different experiments that were performed in triplicates. Statistical significance was determined by Student’s t-test, * p<0.05, ** p<0.01).

### Genetic deletion of N-VEGF using CRISPR-Cas9

To show the significance of endogenous N-VEGF expression on cellular physiology, we took advantage of the CRISPR-Cas9 genome editing system to delete only the coding region of N-VEGF (178 aa in mice) without affecting the translation start site of the “classical” VEGF. To measure VEGF and N-VEGF protein expression levels, we isolated soluble proteins from cytoplasmic and nuclear extracts of ΔN-VEGF cells and control NIH3T3 cells and subjected them to Western blot and immunofluorescence analyses. As clearly demonstrated in Fig. 3A and C, N-VEGF was entirely absent following CRISPR-Cas9 deletion. Hence, hypoxia-induced expression and nuclear mobilization of N-VEGF occurred only in wild type cells. Importantly, N-VEGF deletion did not affect VEGF levels (Fig. 3A).

### Impact of N-VEGF deletion on hypoxia-induced genes

To show the effect of N-VEGF deletion on gene expression levels, RNA-seq analysis was performed on deleted cells and wild type cells before and following hypoxia (see Materials and Methods). A gene expression heatmap focusing on the 3,000 genes modified following hypoxia in wild type versus ΔN-VEGF cells before and after hypoxia is presented in Fig. 4A. The genes are grouped according to expression pattern, similarity or dissimilarity, between the two cell types before or following hypoxia (G1-G6, indicated Fig. 4A, and annotations in Table S2). Most genes (66%) in wild type and ΔN-VEGF cells exhibited similar expression patterns following hypoxia, as indicated in groups 1 and 6. Thus, deletion of N-VEGF led to a partial yet significant cellular response to hypoxia. Interestingly, 34% of the genes induced or repressed by hypoxia showed differential gene expression in ΔN-VEGF compared with control NIH3T3 cells (groups G2-G5). Of note, most of these genes showed differential expression in ΔN-VEGF cells even before hypoxia was induced. Following hypoxia, most of these genes showed only partial elevation or downregulation and did not reach the levels of expression observed in normal cells (groups G2 and G5). A small proportion of the genes displayed opposite expression trends under normoxia as compared to hypoxia in the two cell lines (groups G3 and G4). For example, group G3 showed higher levels of expression in the knockout compared to the wild type under normal conditions, while following hypoxia, expression levels were downregulated in the knockout and elevated in the wild type. This observation shows that N-VEGF deletion had an immense impact on gene expression even before and more massively after hypoxic stress (see annotations in Table S2). Since the main focus of this study was to characterize the nuclear role of N-VEGF, we analyzed the impact of its knockout on the 155 genes modified following N-VEGF translocation to the nucleus (Fig. 4B, NLS-N-VEGF vs. ΔNVEGF). Interestingly, out of the 155 genes modified in NLS-N-VEGF cells, 53% exhibited opposite expression patterns in hypoxia-challenged ΔN-VEGF cells, which is much higher to what is expected by random. Annotations of the genes that were upregulated by nuclear mobilization of N-VEGF and downregulated in N-VEGF-depleted cells included angiogenesis and cell survival pathways (Fig. 4B), including genes associated with negative regulation of apoptotic signaling pathway, such as Aqp1, Msn, Sgk1, Src, Nrp1, and Pik3c2a [16, 21-25]. This suggests that N-VEGF expression protects cells against death elicited by stress signals such as hypoxia.

### N-VEGF deletion leads to increased hypoxia-mediated cell death

To examine if N-VEGF plays a role in defending cells against apoptosis-induced hypoxic stress, the effects of three different yet similar stress signals on ΔN-VEGF vs. wild type cells were compared. More specifically, cells were exposed to hypoxia, CoCl2 [26], and serum deprivation [27], three stressors that mimic lack of blood supply. As a control, we induced Endoplasmic Reticulum stress (ER) [28], which does not mimic lack of blood supply and does not lead to VEGF expression. All stressors resembling lack of blood supply led to statistically significant more apoptosis in ΔN-VEGF cells in comparison to NIH3T3 cells (Fig. 4C, compare second and fourth columns in Hypoxia, CoCl2 and Starvation panels). In contrast, ER stress did not elicit a significant change in the apoptotic index between N-VEGF-deleted cells as compared to their WT counterpart cells (Fig. 4C, Tunicamycin panel). Taken together, N-VEGF is essential for mediation of the anti-apoptotic state in stressed cells to overcome the harsh conditions until oxygen or blood supplies are improved. Overall, N-VEGF has a significant nuclear role on hypoxia-induced genes and physiology, as per the following proposed model (Fig.5). Upon hypoxia, the non-canonical isoform L-VEGF is translated and cleaved to VEGF and N-VEGF. Then, N-VEGF is mobilized to the nucleus and participates in the transcriptional regulation of the hypoxic response and in induction of anti-apoptotic genes. While VEGF is secreted and regulates the hypoxic response in neighboring cells, N-VEGF serves as the autoregulatory arm of VEGF.

## Discussion

Hypoxia is a severe cellular stress that leads to cellular injury and cell death. It leads to the upregulation of expression of gene sets that are involved in pathways aimed at minimizing cellular damage and enabling survival. Among the many rescue pathways activated in response to hypoxia are genes involved in the initiation of angiogenesis, in inhibiting apoptotic pathways, and in the maintenance of cellular structure [29]. This was demonstrated in meta-analysis of hypoxic transcriptomes of many cell types [10]. Our data demonstrated typical changes in the transcriptome profile in the NIH3T3 cell response to hypoxia (Table S2). Likewise, nuclear localization of N-VEGF in various normal as well as cancer cell lines and tissues was also reported [3-6, 30]. Therefore, our model of an autoregulatory role for N-VEGF (Fig. 5) is physiologically relevant and represents its genuine potential.

**Fig. 5.**
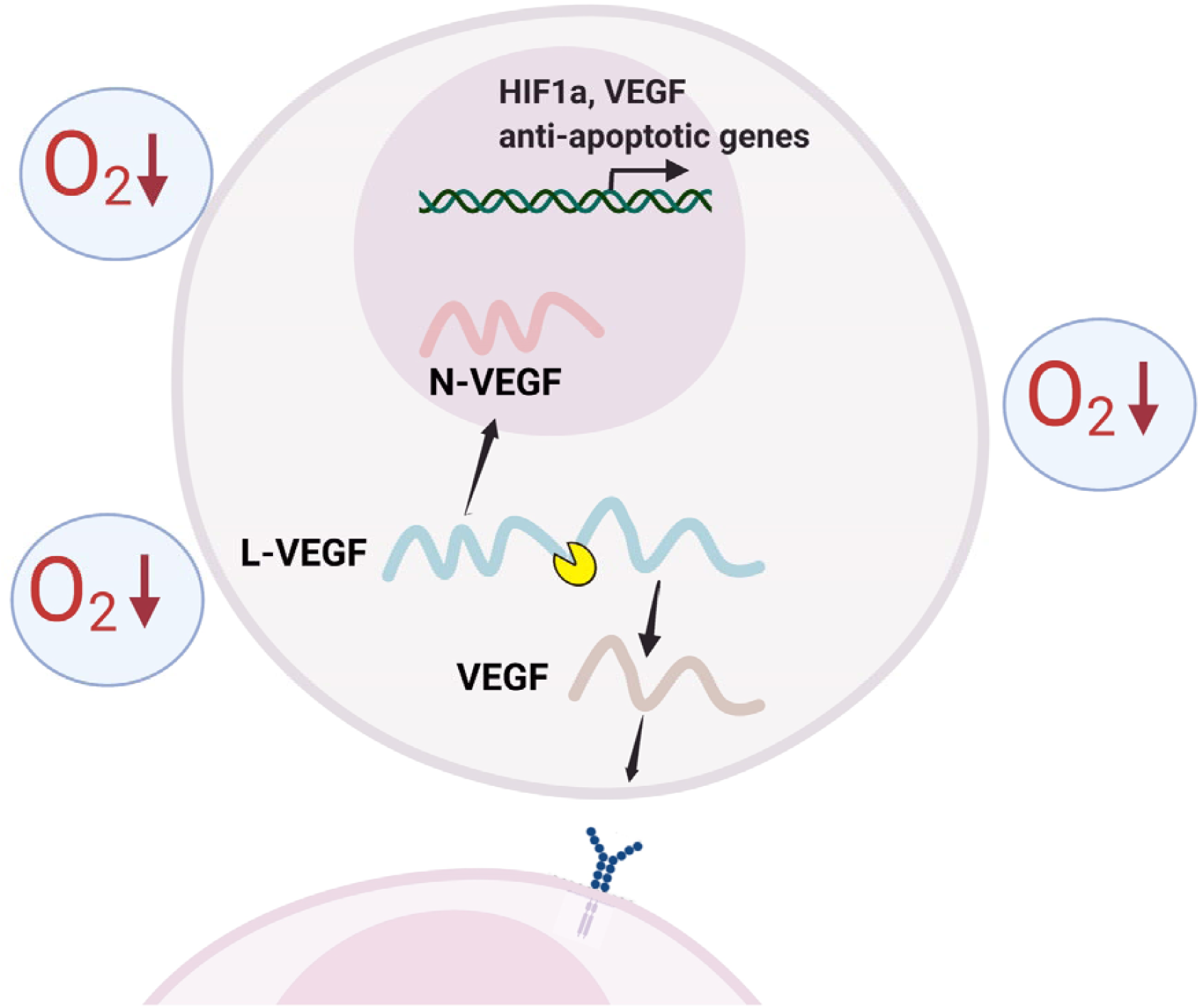
N-VEGF as an autoregulator. Schematic illustration of L-VEGF processing and autoregulation by N-VEGF.

The lack of effect of N-VEGF depletion on VEGF expression levels is explained by the existence of two separate Internal Ribosome Entry Site (IRES) in the 5’ end of VEGF. These two IRES) sequences are functional for non-canonical translation under hypoxic conditions when normal ribosome scanning mechanism is halted (see illustration in Fig. 1A) [9]. N-VEGF deletion results in impaired IRES A and likely IRES B is sufficient for effective translation of VEGF under these conditions [31].

This study provided complementary proof for the essential role of N-VEGF in activating the hypoxic response. We identified an exclusive gene set whose expression was modified by the mere mobilization of N-VEGF to the nucleus. Strikingly, although the hypoxic response is regulated by multiple factors, a significant number of the genes whose expression was modified when N-VEGF was shuttled to the nucleus had the inverse impact when hypoxia was induced in ΔN-VEGF cells. This set of genes included Hif1α and VEGF and a set of genes relevant to hypoxia. Functionally, N-VEGF deletion impaired cell death defense against hypoxic stress.

Although VEGF has been extensively studied in numerous physiological systems over more than three decades and N-VEGF was identified about 20 years ago, our study provides the first evidence of the functional role of N-VEGF. The regulatory architecture of the gene is comprised of two separate arms activated by the same transcript. While VEGF is secreted and activates angiogenesis in surrounding cells, N-VEGF shuttles to the nucleus and regulates intracellular gene responses, thereby serving as the autoregulatory arm of VEGF. This regulatory architecture is beneficial because it provides simultaneous regulation of both intra- and inter-cellular responses by a single promoter (see schematic illustration in Fig. 5). While other biological systems, such as the pro-opiomelanocortin transcript that generates different secreted hormones by proteolytic cleavage (for review see [32]) or the delta-notch signaling in which a segment of the notch receptor is sent to the nucleus and regulates transcription (for review see [33]), to our knowledge, this is the first report of a mechanism in which the same transcript regulates intra- and inter-cellular response. It would be interesting to identify additional transcripts having similar regulatory architecture.

The extent to which N-VEGF plays a regulatory role in hypoxic responses and in angiogenesis in physiological and pathophysiological contexts remains to be determined. Circumstantial evidence suggests involvement of N-VEGF in various pathologies. Genomic polymorphism resulting in IRES B dysfunction was shown to reduce L-VEGF levels in ALS, in macular retinal thickness, and found to be associated with an increased risk of breast cancer aggressiveness, gastric as well as prostate cancers (reviewed in [1]). The role of N-VEGF in models of these diseases as well as in normal development remains to be examined.

Since this study uncovered a previously unknown regulatory mechanism of VEGF, many elementary questions regarding the manner in which N-VEGF exerts its effect remain to be addressed. Future experiments will determine if N-VEGF binds to the chromatin, elucidate the genomic loci it interacts with and identify its partners. In addition, deciphering its three-dimensional structure will help in understanding its specific role.

## Materials and methods

### Cell lines

NIH3T3 (mouse embryo fibroblast) and 293FT (human embryonal kidney) were obtained from ATCC, Manassas, VA (CRL-1658 and CRL-3216, respectively). These cell lines were maintained in DMEM supplemented with 10% fetal calf serum (FCS), amphotericin (2.5 μg/mL) and gentamycin sulfate (50 μg/mL, Biological Industries, Beit-Haemek, Israel).

### Plasmid construction

Lentiviral vector, pLVX-Tet-One-Puro vector (Clontech, #631847), encoding for an engineered Tet activator and the corresponding inducible promoter, which is induced by doxycycline (dox, a derivative of tetracycline), was used. Using PCR a 180 aa N-VEGF containing also 5’ 6xHis-tag was generated, digested with the restriction enzymes EcoRI and AgeI, and cloned into pLVX-Tet-

One-Puro vector termed here N-VEGF. Similarly, a DNA fragment harboring a 5’ 6xHis-tag, an effective NLS, taken from the Cas9 gene and 180aa of N-VEGF was generated by PCR and digested with the restriction enzymes EcoRI and AgeI, and cloned into this vector thus termed NLS-N-VEGF. These clones’ integrity was verified by DNA sequencing.

### Lentiviral production

4×10^6^ 293FT cells were seeded in 10cm tissue culture plate 24hrs prior to transfection. Five µg of lentiviral vector, 1 µg pMD.G and 4 µg psPAX2 were transfected using CalFectin, mammalian transfection reagent (SignaGen, Maryland, USA) according to manufacturer’s instructions. Twenty-four hrs post-transfection medium was changed with 7 ml of fresh medium. Forty-eight hrs post-transfection viral supernatant was collected, filtered using 0.2µm syringe-filter and aliquots were flash-freezed using liquid nitrogen. The viral supernatant was stored at −80 ^0^C for further use.

### NIH3T3 infection

Twenty-four hrs prior to infection, 2.5x 10^5^ cells were seeded in a 12 well tissue plate with 0.5 ml of medium. Then, 0.5 ml of viral supernatant and 8 µg/ml of polybrene were added to each well. The cells were incubated at 37^0^C in 5% CO_2_ incubator. Twenty-four hrs post-transfection, the cells were split 1:2 into 6 well tissue culture plates. Stably infected cells were selected with Puromycin (3µg/ml) for 48 hrs post-infection and resistant cells were harvested 96 hrs later.

### Nuclear and cytoplasmic fractionation

NIH3T3 cells were trypsinized (2×10cm plates) and washed twice with ice cold PBS and lysed on ice for 15min in 100µl of cytoplasmic lysis buffer (10mM HEPES pH 7.4, 10mM KCl, 0.01mM EDTA, 0.1mM EGTA, 2mM DTT, 5mM Na¬2VO4, 0.1% NP40 and cOmplete, EDTA-free protease inhibitor cocktail (Sigma-Aldrich). Nuclei were pelleted using benchtop microfuge (6,000rpm for 2 min) and the supernatant containing the cytoplasmic fraction removed to which Urea and SDS were added to a final concentration of 2M and 2%, respectively. The samples were denatured by boiling for 5 min. The nuclei were then washed twice in 1ml of cytoplasmic lysis buffer to remove any cytoplasmic contaminant, and re-sedimented. Subsequently, the nuclei were lysed in 100μl of denaturing lysis buffer (2% SDS, 2M urea, 8% sucrose, 1mM NaF, and 5mM Na¬2VO4). Genomic DNA was sheared by passage through a Qiashredder column (Qiagen, Crawley, UK) in a benchtop microfuge (6,000rpm for 2 min) and denatured by boiling for 5 min.

### SDS-PAGE and Western blotting

SDS-PAGE and Western blotting were preformed using Bio-Rad Mini-protein apparatus. Gel was prepared and blotted, and reacted with antibodies exactly as previously described [6]. The primary antibody used was Rabbit polyclonal αORF (N-VEGF), made in our laboratory, that was affinity purified against 5’UTRORF of VEGF (1:500 dilution in TBST+2.5% non-fat dried milk-powder (Carnation)). The primary antibody was incubated for 3 hrs at R.T or O.N. at 4^0^C. Secondary antibody, HRP anti-rabbit (1:10,000 dilution in TBST+2.5% non-fat dried milk-powder) was incubated for 1-2 hrs. Bands were visualized using ECL2 Western Blotting kit (Thermo Scientific, Waltham, MA).

To demonstrate even loading of protein samples, membranes were stained, prior to membranes blocking, with Ponceau-S (Sigma-Aldrich) according to the manufacturer’s instructions and exactly as described before [34]. The purity of cytoplasmic and nuclear soluble protein extracts was determined by Western blot. Membranes were striped from anti N-VEGF or VEGF polyclonal antibodies and following re-blocking reacted with primary antibodies directed against Lactate Dehydrogenase (anti-LDH) (Abcam, cat #ab47010,1:300) and secondary antibodies anti-goat (1:10,000), to test nuclear extract purity. Subsequently the membrane was striped once more from bound antibodies, re-blocked, and reacted with antibodies directed against Histone H3 acetyl K27 (αH3K27ac) primary antibody (Abcam, #ab4729, 1:500) and secondary antibodies, anti-rabbit (1:10,000) to test cytoplasmic extract purity.

### Immunofluorescence staining and microscopy

NIH3T3 cells, transfected with a vector driving the expression of either N-VEGF or NLS-N-VEGF, were plated in 24 wells plates containing microscope coverslips 0.13 mm. Some of the samples were treated with 1 µg/ml dox for 48 hrs. After additional 24 hrs, the cells were either not treated or subjected to hypoxia for additional 16 hrs. Cells were fixed with 4% PFA for 15 min at room temperature, washed two times with PBST, permeabilized with 0.1% Triton X-100 for 15 min and washed three times. Cells were then blocked with 5% normal goat serum in TBST for 1 hrs and subjected to staining with primary antibodies directed against N-VEGF (1:100 dilution in blocking solution) for O.N at 4 ^0^C in a humidified chamber with very gentle swirling. Finally, the coverslips were washed four times with PBS. Slides were subsequently reacted with fluorescent secondary antibodies Alexaflour 488 goat anti rabbit (H+L) (1:1000 dilution in blocking solution) for 1 hr in the dark at RT. Following incubation, the slides were washed three times with PBS and reacted with DAPI and Fluromount-G for nuclei staining. Immunostained cells were visualized under Zeiss LSM 700 laser scanning confocal microscope, Green laser – 555 nm (10mW), Red laser – 639 nm (5mW), magnification of X30 for N-VEGF transfected cells and magnification of X60 for NLS-N-VEGF transfected cells (Fig 1C). Alternatively, cells were visualized under Zeiss Cell Observer inverted fluorescence microscope, magnification x20 (Fig 4D).

### Real-time RT-PCR

The primers used for real-time PCR were designed using PrimerExpress™ software (Applied Biosystems; see Supplemental Table S3). 800 ng total RNA were reverse transcribed to cDNA using High-Capacity cDNA Reverse Transcriptase kit (Ambion) according to manufacturer’s instructions. cDNA was amplified with two primers for each gene using Power SYBR Green PCR Master Mix (Applied Biosystems, USA) and Real-time PCR analysis was performed using QuantStudio 12K Flex (Applied Biosystems by Life technology, USA), according to manufacturer’s instructions. Only primer pairs exhibiting PCR efficiency of 90% or higher were chosen. Gene expression was normalized to GAPDH mRNA expression. The data is presented as the relative expression of the gene of interest compared with GAPDH. The mRNA expression trend of genes mentioned in this publication (up, down or unchanged) that were determined by RNA-seq, were all validated by independent real time RT-PCR.

### RNA-seq

RNA samples concentration and integrity were measured using Qubit (ThermoFisher) and TapeStation (Agilent), respectively. Libraries for sequencing were prepared using the TruSeq RNA Library Prep Kit v2 (Illumina) according to manufacturer’s instructions. Libraries were sequenced on Illumina NextSeq 550 instrument with 80 bps single-read. The number of reads ranged from 54,915,873 to 38,372,655 per sample. Library quality control was conducted using FASTQC, version 0.11.5. Sequences were trimmed by quality using trim galore and aligned to Mus musculus genome build – GRCm38.p6 with release 94 annotation file using Tophat2, version 2.1.0 (uses Bowtie2 version 2.2.6). Raw gene expression levels were counted by HTseq-count, version 0.6.1. All downstream analyses were conducted in R with DESeq2 [35] as follows: raw data were filtered by independent filtering. Count normalization was performed by DESeq2 default. To identify differentially expressed genes differing between two conditions we used the Wald test, as implemented in DESeq2.

Heatmaps were plotted using R pheatmap package. For the heatmap normalized counts were scaled by row and the genes were ordered as specified in the figure legends.

All RNA sequencing data were deposited to the Gene Expression Omnibus (GEO) with accession numbers GSE178297 and GSE178298.

### CRISPR/Cas9-mediated deletion of N-VEGF

The strategy for generating N-VEGF deleted cells is shown in Fig. S4. sgRNAs from the 5′ and 3′ ends of the 180 aa long N-VEGF were designed using the chopchop online tool [36], and highly scored sgRNAs were chosen (sgRNA 1–6). Each sgRNA was phosphorylated, annealed, and cloned to lentiGuide-Puro (addgene no. 52963), which was previously digested with BsmbI. In order to create gRNAs with sticky ends complementary to BsmbI digestion product, “CACC” and “AAAC” were added to 5′ end of the designed 20-bps sgRNA in the forward and reverse oligomers, respectively. In order to verify the efficiency and specificity of the sgRNAs, a T7 endonuclease assay was performed on NIH3T3 cells and the cleavage efficiency was analyzed using TapeStation (Agilent). Two gRNAs (1 and 4) were found to specifically targeted the ORF: gRNA1 target-site PCR amplicon (500 bps) was cleaved to 300 and 200 bp fragments, and gRNA4 target-site PCR amplicon (461 bp) was cleaved into 329 and 132 bp fragments (data not shown). Briefly, a two steps protocol was utilized to generate NIH3T3 cells with biallelic deletion of the ORF. To get biallelic clones with reporter cassette insertion with high probability, cells were transfected with the two gRNAs and Cas9 encoding vector and two donor DNA segments (with different selectable markers, Puromycin and Hygromycin) and cultivated for 2 weeks. Single clones, resistant for the two antibiotics and containing mCherry reporter gene instead of the N-VEGF sequence, were picked and the 5′ and 3′ junctions were assayed by PCR (Fig. S5). Four Clones harboring the biallelic N-VEGF replacement cassette were further verified by DNA sequencing and taken for further study. For “clean” deletion of the N-VEGF two clones were transduced with pMSCV-VCre encoding retroviral vector [37] to remove the reporter\selection cassettes (Fig. S6). Three days later, non-fluorescent cells were harvested following two cycles of FACS sorting. N-VEGF deletion was verified by PCR and by DNA sequence analysis.

### Flow cytometry

Flow cytometry sorting was performed using BD FACSAria-IIIu cell sorter (BD Bioscience), and data was analyzed using Flowing Software 2 (Cell Imaging Core, Turku Centre for Biotechnology). Unstained WT NIH3T3 cells were used as negative control and NIH3T3 cells containing mCherry reporter gene were used as positive control.

### Hypoxia treatment

Hypoxic cultures were maintained in a hypoxia chamber (<0.2% O2) for 16 hrs at 37°C in humidified 95% N2/5% CO2, and normoxic cultures were maintained in 95% air/5% CO2 for similar duration.

### ApoTox-Glo™Triplex assay

BD BioCoat Poly-D-Lysine 96-well plates with black sides and clear bottoms were used to run the ApoTox-Glo™ Triplex Assay (Promega). Cells were trypsinized and removed from their culture flasks and diluted to 24,000 cells/ml and 100µl of each sample was added to each well. Viability and Cytotoxicity reagents from the kit were prepared according to manufacturer’s instructions, added to a 96-well plates, and the plates was incubated at 37°C for 1 hr. After incubation, plates were analyzed for fluorescence as defined in the protocol using a Spark® multimode microplate reader by Tecan. The caspase reagents were then added to the plates and the plates were incubated at RT for an additional 1 hr. Luminescence was then measured.

### Cobalt chloride and Tunicamycin treatments

Cobalt chloride (Spectrum) was added to a final concentration of 550 µM to N-VEGF NIH3T3 and the counterpart part ΔN-VEGF cell cultures. The live and dead cells were counted using trypan blue staining after 4, 6, 8 and 17 hrs of treatment. Next, the ApoTox-Glo™Triplex assay was used to test the viability, cytotoxicity, and apoptosis of the cells as detailed above.

Tunicamycin (Sigma) was added to a final concentration of 0.5 µg/ml to N-VEGF NIH3T3 and the counterpart part ΔN-VEGF cell cultures. The live and dead cells were counted using trypan blue staining after 5 hrs of treatment. Then the ApoTox-Glo™Triplex assay was used to test the viability, cytotoxicity, and apoptosis of the cells.

### Statistical analysis

Each experiment was conducted at least three times in triplicates unless otherwise stated. Values are presented as means±AvDev. Except for Fig. 10, the data was compared by unpaired two-tailed Student’s t-test; p-values <0.05 (*), <0.01(**) or <0.001 (***) were considered as statistically significant, as indicated in the appropriate figure. As indicated in some figure legends, two-way ANOVA was performed on the differences with cell type and various stress stimuli as main factors. The ANOVA was followed by post-hoc analysis of all possible contrasts using Tukey’s honestly significant difference, which controls for multiple comparisons. Contrasts with p-values <0.05 are considered significant. The statistical analyses were performed using the R language and environment for statistical computing and graphics.

## Acknowledgements

We are grateful to the Gellman-Lasser Fund for Medical & Biomedical Research and innovation (grant No. 907679) to B.Z.L, and to the Technion’s Genome Center and the Russell Berrie Nanotechnology Institute for their support. We are also deeply grateful to Prof. Gera Neufeld, Faculty of Medicine, Prof. Yuval Shoham, Faculty of Biotechnology and Food engineering and Dr. Nitsan Fourier, Genomic Centre, Technion, for their input and critical reading of the manuscript. We would like to thank Dr. Yehuda Shabtai from UPenn for fruitful discussions.

The initial construction of the Tet-On lenti-vectors by Mrs Tania Vorobyov, research assistant, is acknowledged. B.Z.L. is an incumbent of the Lily and Silvian Marcus Chair in Life Sciences, Technion.

## Supporting Information

**Fig. S1.**
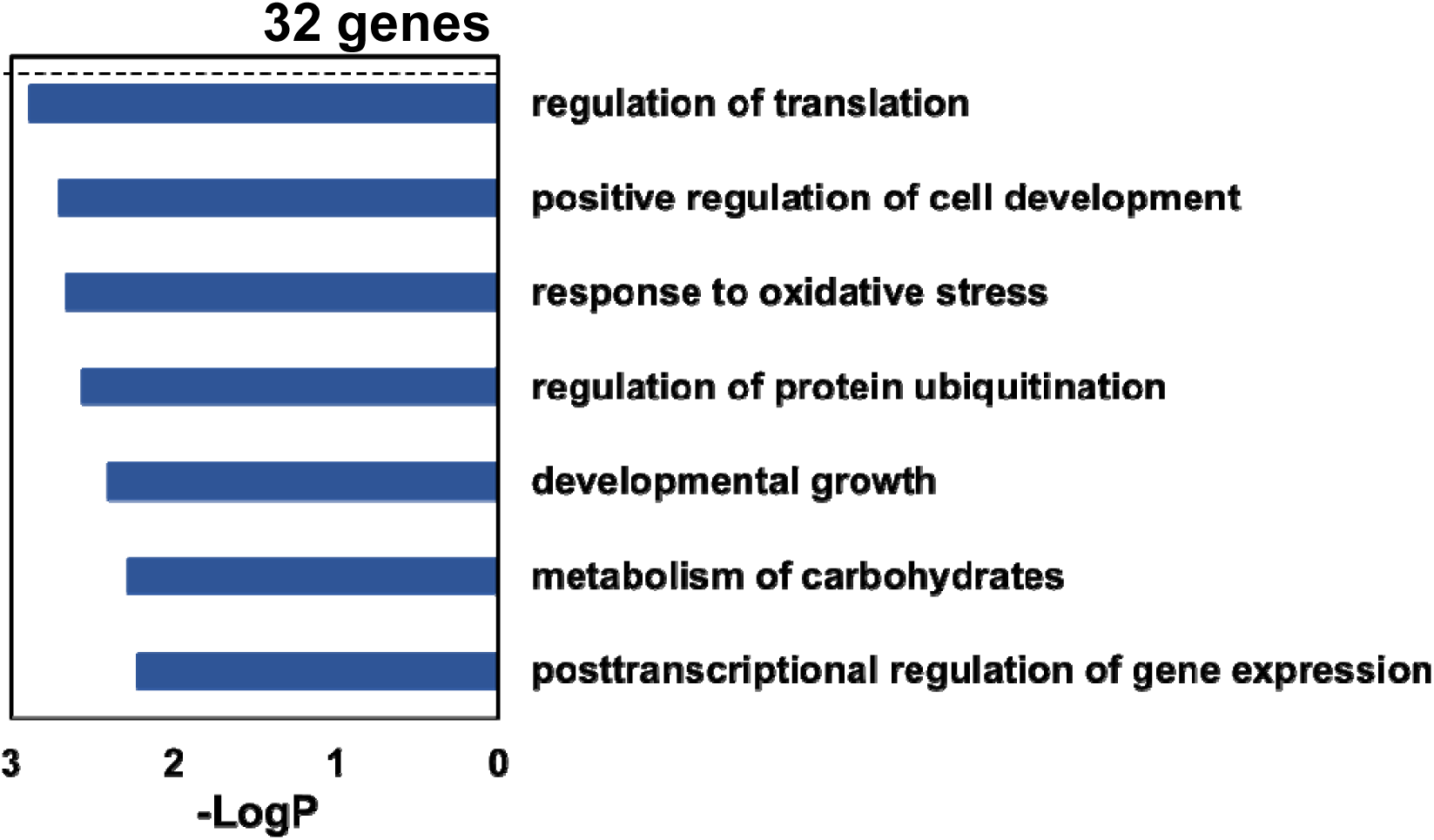
Annotation and pathway analysis of the 32 differentially expressed genes that were down regulated following hypoxia independent nuclear mobilization of NLS-N-VEGF. The annotations for the downregulated genes was performed as elaborated under Fig.1.

**Fig. S2.**
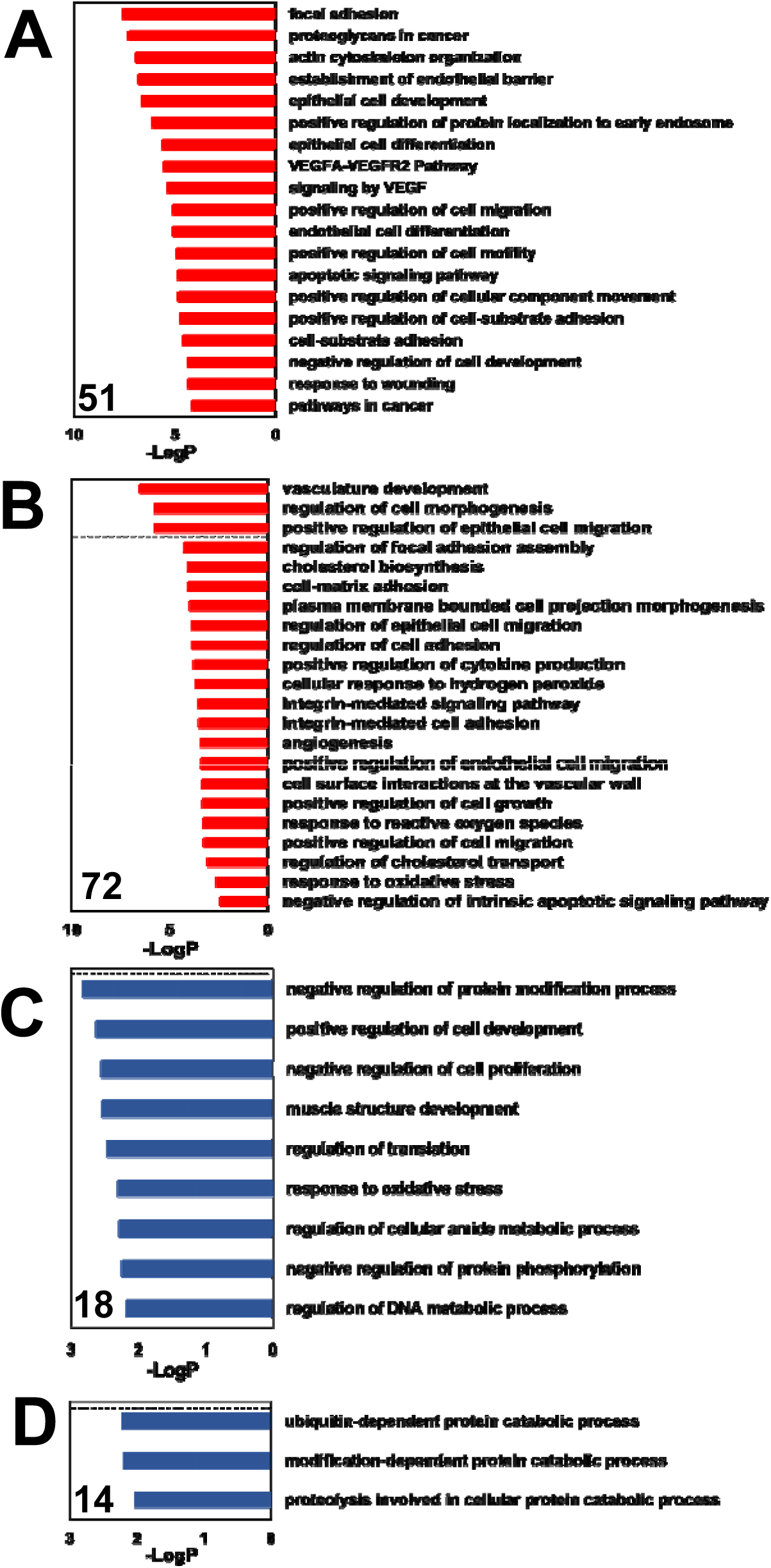
Annotation of 155 genes that exhibited differential expression in NLS-N-VEGF cells and NIH3T3 cells following hypoxia. (A) Annotation of genes (51) that were upregulated in both cell types. (B) annotation of genes (72) that were upregulated in NLS-N-VEGF cells following dox and not changed or downregulated in NIH3T3 cells. (C) annotation of genes (18) that were downregulated in NLS-N-VEGF cells following dox and upregulated or not changed in NIH3T3 cells. (D) Annotation of genes (14) that were downregulated in both cell types. Bars below dotted line represent less significant annotation groups.

**Fig. S3.**
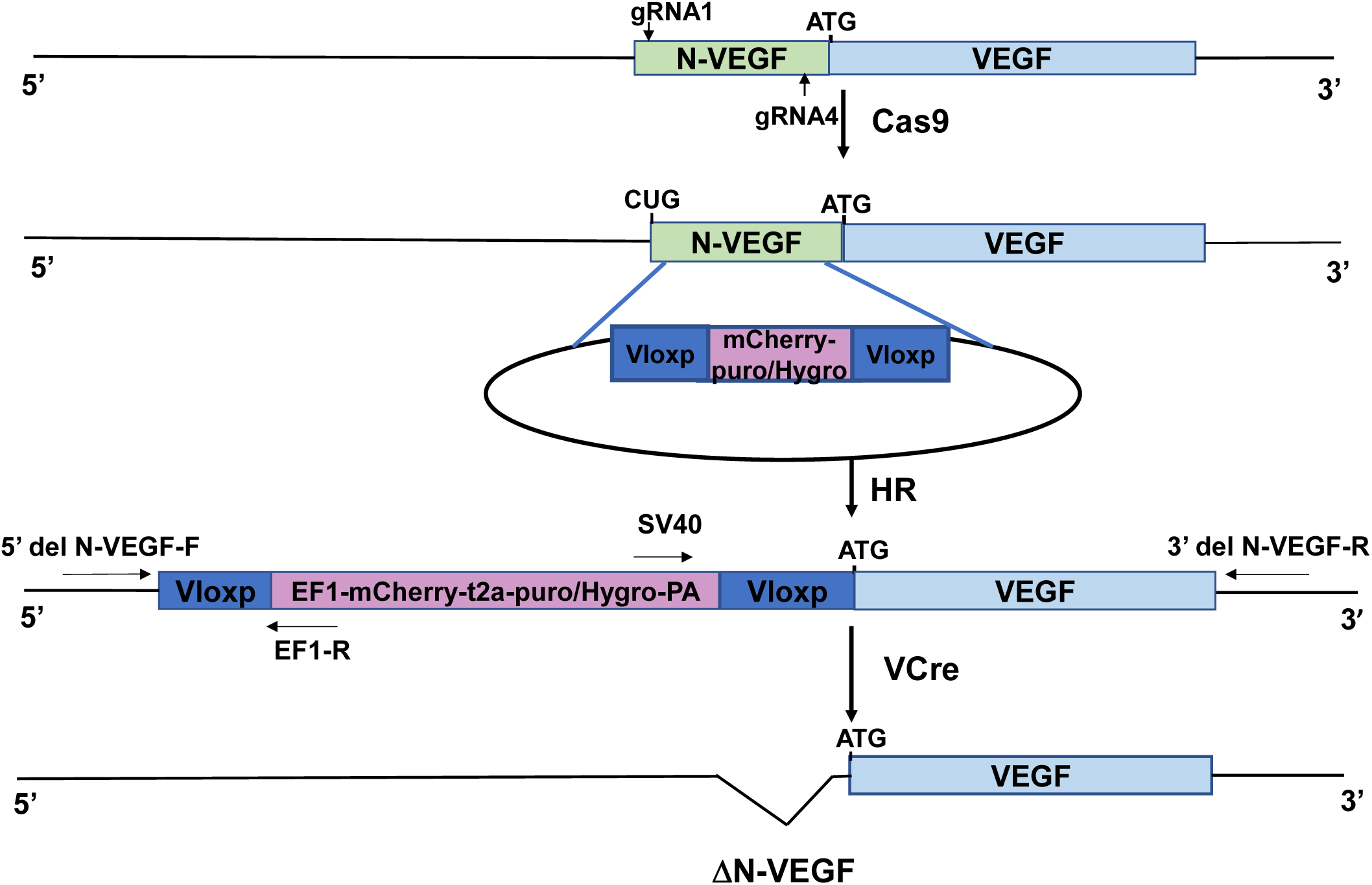
Schematic illustrations. Schematic illustration of the two-steps protocol for CRISP-Cas9 mediated biallelic deletion of N-VEGF.

**Fig. S4.**
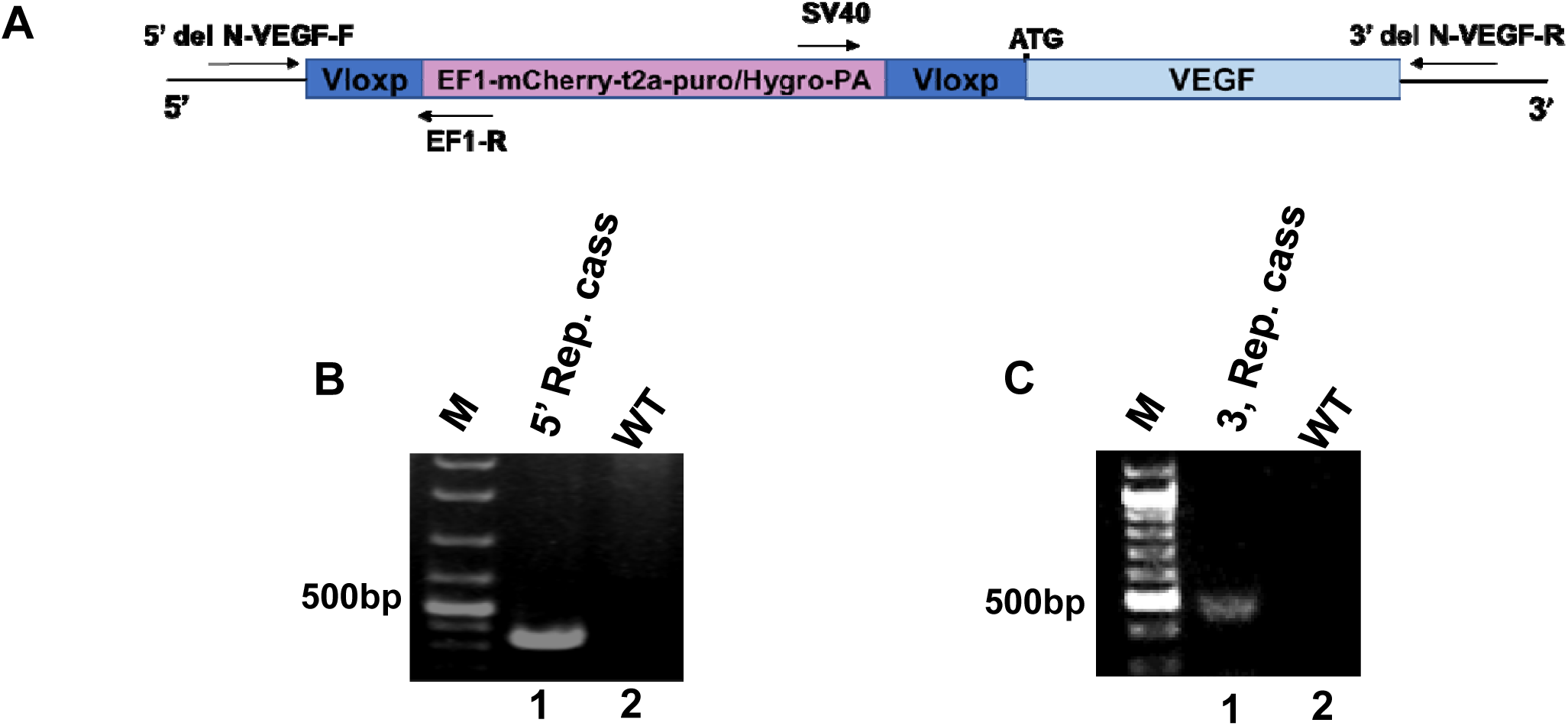
PCR analysis of NIH3T3 with a replacement of the N-VEGF gene with a reporter cassette. **(A)** Schematic illustration of the reporter cassette harboring a reporter gene followed by Puromycin or Hygromycin resistance genes. Primer pairs for each junction were used for clones analysis. **(B)** Agarose gel showing the expected 435bp PCR product from 5’ junction of isolated clone 1 (panel 1) obtained using the 5’-del NVEGF forward primer and reverse primer EF1-R from the insert. **(C)** Agarose gel showing levels of the expected 482bp PCR product from the 3’ junction (panel 1), obtained using the SV40 forward primer and 3’ del NVEGF reverse primer.

**Fig. S5.**
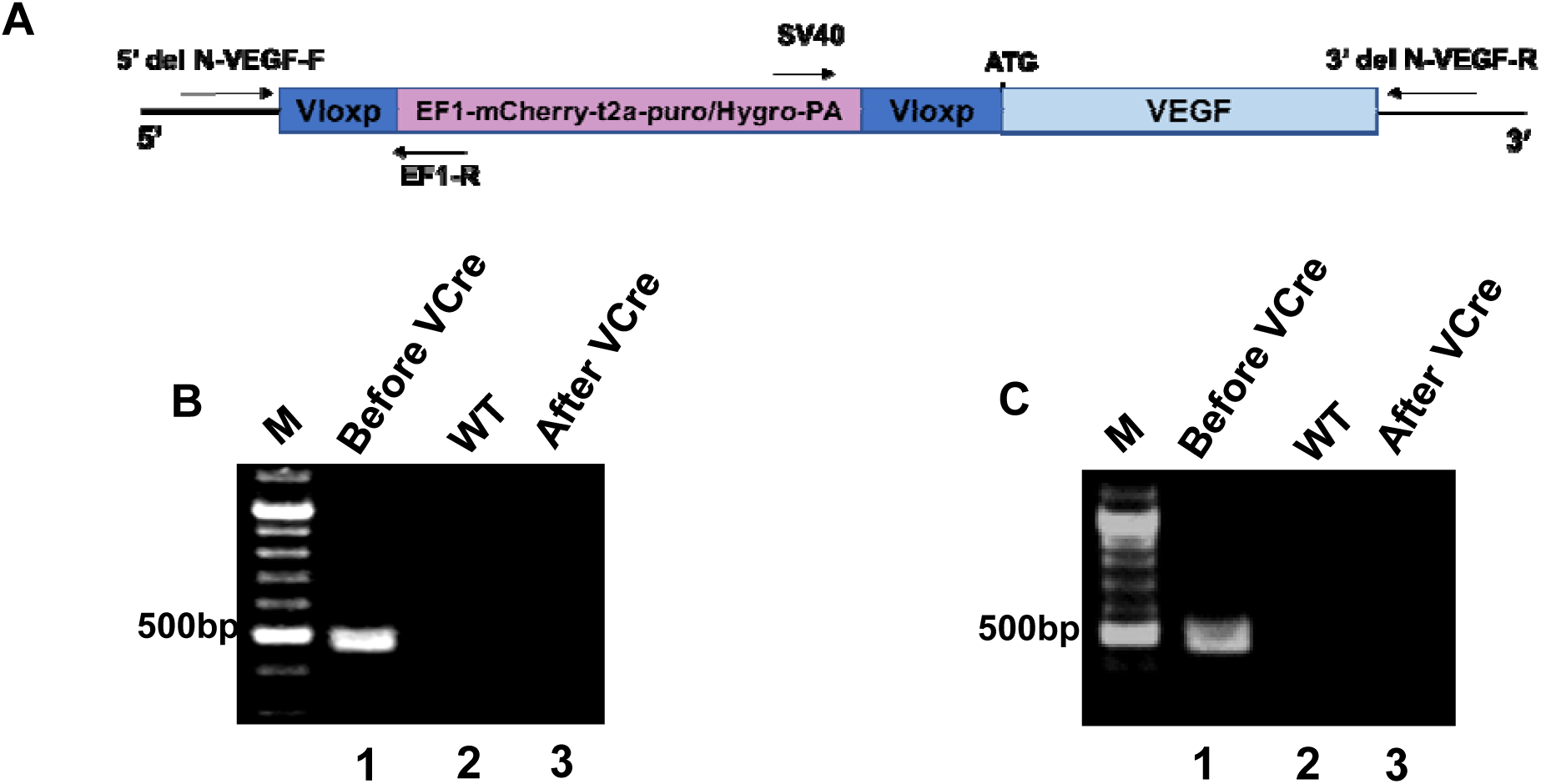
PCR analysis to verify genomic N-VEGF deletion from NIH3T3. (A) Schematic illustration of the reporter cassette harboring a reporter gene followed by Puromycin or Hygromycin resistance genes. Primer pairs for each junction were used for clones analysis. **(B)** Agarose gel showing levels of the expected band of 435 bp PCR product from the 5’ junction of clone 1 before (panel 1) and following VCre expression (panel 3). Removal of the reporter cassette was determined using 5’-del NVEGF forward primer and reverse primer EF1-R from the inset. **(C)** Agarose gel showing levels of the expected band of 482 bp PCR product from the 3’ junction, obtained using the SV40 forward primer and 3’-del NVEGF reverse primer before (panel 1) versus after VCre expression (panel 3).

**Table S1. Gene list and annotation of 155 genes the exhibited differential expression in NLS-N-VEGF cells treated with dox**. Annotation of 123 upregulated genes (up tab) and 32 downregulated genes (down tab) in NLS-N-VEGF cells treated with dox.

Table S1.xlsx

**Table S2. Annotation of genes that exhibited differential expression between Δ-N-VEGF and NIH3T3 cell following hypoxia**. The genes are arranged in six groups (G1-G6). G1 and G6, similar expression pattern in both cell types of genes either induced or repressed following hypoxia, respectively. G3 and G4, inverse expression pattern in Δ-N-VEGF cells, either upregulated (G3) or downregulated (G4) following hypoxia in comparison to NIH3T3 under the same treatment. Last, genes that exhibited no change in expression pattern in ΔN-VEGF cells that were either upregulated or downregulated in NIH3T3 cells following hypoxia, groups G2 and G5, respectively. Statistically significant annotation groups, Log q < 0.05.

Table S2.xlsx

**Table S3. List of primers used in this study**.

Table S3.xlsx

## Notes

### Competing Interest Statement

The authors have declared no competing interest.

### Summary of Updates

Vascular endothelial growth factor A (VEGF-A) is a secreted protein that stimulates angiogenesis in response to hypoxia. Under hypoxic conditions, a non-canonical long isoform called L-VEGF is concomitantly expressed with VEGF-A. Once translated, L-VEGF it is proteolytically cleaved to generate N-VEGF and VEGF-A. Interestingly, while VEGF-A is secreted and affects the surrounding cells, N-VEGF is mobilized to the nucleus. This suggests that N-VEGF participates in transcriptional response to hypoxia. In this study, we performed a series of complementary experiments to examine the functional role of N-VEGF. Strikingly, we found that the mere expression of N-VEGF followed by its hypoxia-independent mobilization to the nucleus was sufficient to induce key genes associated with angiogenesis, such as Hif1alpha, VEGF-A isoforms, as well as genes associated with cell survival under hypoxia. Complementarily, when N-VEGF was genetically depleted, key hypoxia-induced genes were downregulated and cells were significantly susceptible to hypoxia-mediated apoptosis. This is the first reports of N-VEGF serving as an autoregulatory arm of VEGF-A. Further experiments will be needed to determine the role of N-VEGF in cancer and embryogenesis.

